# Social groups buffer maternal loss in mountain gorillas

**DOI:** 10.1101/2020.09.01.276956

**Authors:** Robin E Morrison, Winnie Eckardt, Fernando Colchero, Veronica Vecellio, Tara S Stoinski

**Affiliations:** Dian Fossey Gorilla Fund, Rwanda; Centre for Research in Animal Behaviour, University of Exeter; Department of Mathematics and Computer Science, University of Southern Denmark; Interdisciplinary Center on Population Dynamics, University of Southern Denmark

## Abstract

Mothers are crucial for mammals’ survival before nutritional independence, but many social mammals reside with their mothers long after. In these species the social adversity caused by maternal loss later in life can dramatically reduce fitness. However, in some human populations these negative consequences appear to be overcome by care from other group members. We investigated the consequences of maternal loss in mountain gorillas and found no discernible fitness costs to maternal loss through survival, age at first birth or survival of first offspring through infancy. Social network analysis revealed that maternal loss led to strengthened relationships with other group members, particularly the dominant male and age-mates. In contrast to most social mammals, where maternal loss causes considerable social adversity, in mountain gorillas, as in certain human populations, this may be buffered by relationships within cohesive social groups, breaking the link between maternal loss, increased social adversity and decreased fitness.

## Introduction

Maternal loss, along with a number of other indicators of early-life adversity, is one of the strongest predictors of lifespan in humans and other social mammals^1^. In mammals, mothers are vital for the survival of young offspring which are nutritionally dependent on their mother’s milk. In some species, particularly social species with slow life histories^2^, mothers continue to provide benefits to their offspring throughout immaturity and even into adulthood and can dramatically influence their fitness. In social mammals the removal of maternal care due to maternal loss can reduce survival in the youngest offspring^3^ by affecting for example nutrition, thermoregulation and protection^4^. But maternal loss can also reduce the fitness of offspring in a far broader age range through long-term effects on their social environment, negatively influencing their social integration and social status throughout their lives^1,5,6^. Multiple studies have now confirmed effects on survivorship for individuals orphaned well past the period of nutritional dependency, with these negative changes to their social environment, often termed social adversity, posited to be the key mechanism by which this occurs^5,7–10^. In social mammals, the social environment can have extreme consequences for health, fitness and lifespan, mediated through pathways such as chronic stress, immune function or environmental exposure^1^. In humans, an individual’s social environment, specifically, the strength of their social relationships, predicts a ∼50% increase in human mortality, comparable to the effects of smoking and alcohol consumption, and exceeding those of physical inactivity and obesity^11^.

The impact of maternal loss varies based on the loss in benefits relative to individuals with mothers present. In species with sex-biased dispersal, the consequences of maternal loss can differ between the sexes due to longer periods for potential investment in non-dispersing offspring^4,12–14^. In female-philopatric red deer, maternal loss increases mortality for males and females, but this effect is only detectable in males <2 years, whilst for females maternal benefits continue throughout their lives^7^. In male-philopatric chimpanzees, males that suffer maternal loss before reaching 15 have lower survival, whilst females only show reduced survival from maternal loss under the age of 10^8,15^. In killer whales, where neither sex disperses, maternal loss when offspring are >30 years old reduces survival for both sexes^10^. However, this reduction is considerably greater for males and maternal loss between 15 and 30 years appears to reduce male but not female survival. This is thought to be due to higher maternal investment in males which mate outside the group and who’s offspring therefore don’t increase within-group feeding competition.

Several studies have demonstrated the specific social benefits that mothers provide to co-residing offspring, which can extend well into adulthood^7,8,16,17^. In some societies the active support of mothers can increase the rank of their offspring^6,18–20^ with higher ranking individuals showing greater survival in almost all natural animal populations in which it has been studied^1^. In baboons, where female rank is inherited from the mother, mothers play a key role in integrating their offspring within the larger social group^5^. Maternal loss in baboons leads to lower social integration^5^, with less integrated individuals suffering higher mortality^21^. Maternal presence can also influence nutrition. Chimpanzees that suffer maternal loss up to late juvenility have significantly lower muscle mass^22^. This was suggested to be due to mothers buffering against feeding competition and increasing juvenile’s access to valuable resources. Mothers can also provide valuable ecological knowledge either directly or through their relationships with more experienced group members^8^. African elephant groups led by more experienced matriarchs showed greater foraging success during extreme drought^23^, whilst in resident killer whales, older females led salmon foraging, particularly when abundance was low^24^. Mothers can also benefit offspring by increasing opportunities for social learning of complex feeding techniques such as termite-fishing^25^ or nut-cracking in chimpanzees, where immatures nut-cracking efficiency correlates positively with that of their mothers^26^.

As a result of these numerous benefits, it is thus not surprising that maternal loss not only influences offspring survival but can impact other components of their offspring’s fitness, such as reproduction and the survival of grand-offspring. Male bonobos residing in groups with their mothers sire three times the number of offspring^16^, whilst maternal loss before weaning negatively affects antler development in male red deer – a trait found to correlate with reproductive success^7^. In chimpanzees, females mature faster, first give birth younger^27^ and enter the dominance hierarchy higher^28^ if their mothers are present which is expected to considerably increase their lifetime reproductive success. In baboons, if mothers had themselves suffered maternal loss in the first 4 years of their life, their offspring had 48% higher mortality throughout the first 4 years of their life, suggesting an intergenerational effect of maternal loss driven by lifelong developmental constraints^29^. Maternal loss in social mammals with extended maternal care can therefore have long-term fitness consequences mediated through multiple pathways that detrimentally affect survival and reproduction.

Due to the extended periods of mother-offspring co-residence and the important social support mothers can provide, highly social species often have the most to lose from maternal loss. However, social groups also provide the potential for support from other group members following maternal loss. Both kin and non-kin group members have been suggested to compensate for the loss of close kin to varying extents^30–33^ and the strengthening of relationships with remaining group members may buffer against changing social environments^34^. In chacma baboons social support from group members is thought to alleviate the stress of losing a close relative^31^. Similar social support has been suggested in elephants which associate more with age-mates and siblings in response to maternal loss^32^. However, these orphaned elephants interact less with matriarchs which may decrease their access to key knowledge and high-quality resource patches. In chimpanzees, older siblings can ‘adopt’ younger siblings after maternal loss, increasing their social contact and showing heightened vigilance in dangerous situations^33,35^. But despite these compensatory social behaviors, the negative consequences of maternal loss post weaning are well documented in all three genera^5,8,15,22,32,36^.

In humans, the death of a mother was associated with increased child mortality in all 28 historic and contemporary populations studied by Sear and Mace^37^. However, this effect appeared to decline substantially with age, disappearing for children that suffered maternal loss >2 years in 5 of the 11 populations in which it was investigated^37^. This reduced mortality is thought to be due to the care provided by other kin, particularly after weaning, suggesting that social buffering from other group members can overcome the negative effect of maternal loss on survival in certain circumstances. Whilst the effects of care from specific kin members varied across populations, at least one kin member significantly impacted child survival in all studies^37^, and there is evidence for the importance of maternal grandmothers^38,39^ and fathers^40,41^ in particular. Killer whale maternal grandmothers, especially those that are post-reproductive are also known to improve offspring survival^42^. In humans, there is also evidence for the benefits of care provided by non-kin such as step-mothers^43,44^ and through the modern practices of non-kin adoption^45^.

Mountain gorillas (*Gorilla beringei beringei*) show extended maternal care with offspring remaining in their natal groups until at least sexual maturity and approximately half remaining beyond sexual maturity (48% of females^46^ and 55% of males^47^). Females that disperse from their natal group tend to do so earlier (mean age of 7.9 years^46^) than males (mean age of 15.3 years^47^) and therefore have a shorter period of potential maternal investment. The complexity of gorilla social structure suggests that detrimental long-term effects on their social environments could have particularly negative fitness consequences^48–50^. However, these stable, cohesive, social groups also have the potential to provide a social buffer to the negative consequences of maternal loss. Mountain gorilla groups either contain a single adult male (approximately 64% of groups), or multiple adult males (approximately 36% of groups)^51^, at least one adult female, and their offspring^52^. Single male groups are polygynous whilst multimale groups have high reproductive skew toward the dominant male who sires the majority of offspring^53–55^. Infants (<4 years) are nutritionally dependent on their mothers until being weaned at a mean of 3.3 years^56^ and are reliant on their mothers for thermoregulation and transport, being carried for prolonged periods^57^. Juveniles (4-6 years) are nutritionally independent but remain in close proximity to their mothers the majority of the time^57^. Dominant males’ primary form of care is through protection from out-group males and potential predators^58^. But they also show high levels of affiliative behaviour towards infants, grooming and resting in contact with them, with no evidence that they discriminate between infants based on paternity^59,60^.

In this study, we use the long-term demographic records of the Dian Fossey Gorilla Fund’s Karisoke Research Center collected over 53-years (1967 – 2019) to investigate the potential for social relationships to buffer the negative social consequences of maternal loss in mountain gorillas. We quantify the effects of maternal loss on multiple fitness measures: survival, female age at first birth, female survival of first offspring through infancy and male dominance. We then use social proximity and contact data collected between 2003 and 2015 to investigate the social responses of group members to maternal loss by 31 immature gorillas, and the potential for social buffering by group members to compensate for maternal loss.

## Results

### Effect of maternal loss on survival

To determine the effect of maternal loss on survival, we carried out a Cox-proportional hazards analysis separating individuals based on four maternal orphan (hereafter, orphan) classes based on the age at which their mother died: a) infants (2-4 years), b) juveniles (4-6 years), c) sub-adults (6-8 years) and d) non-orphans (>8 years) if mothers died after reaching maturity. Due to the small sample sizes for the juvenile and sub-adult classes, analysis was also run with these two classes merged. We found no significant differences in survival between all orphan classes and the non-orphan class for both sexes irrespective of using three or four classes (Table 1, Figure 1, Supp. Table 1).

**Table 1.**
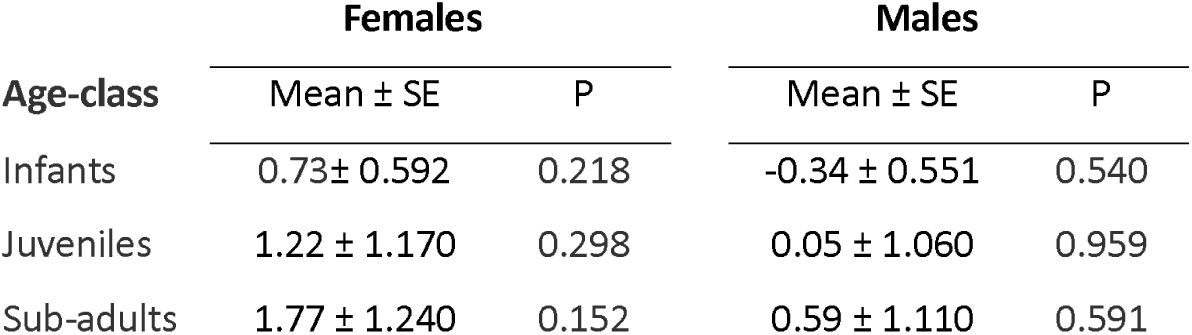
Cox-proportional hazards models showing the effects of the four age-classes on survival. All results are relative to the non-orphan class.

| Age-class | Females |  | Males |  |
| --- | --- | --- | --- | --- |
| | Mean ± SE | P | Mean ± SE | P |
| Infants | 0.73 ± 0.592 | 0.218 | -0.34 ± 0.551 | 0.540 |
| Juveniles | 1.22 ± 1.170 | 0.298 | 0.05 ± 1.060 | 0.959 |
| Sub-adults | 1.77 ± 1.240 | 0.152 | 0.59 ± 1.110 | 0.591 |

**Figure 1.**
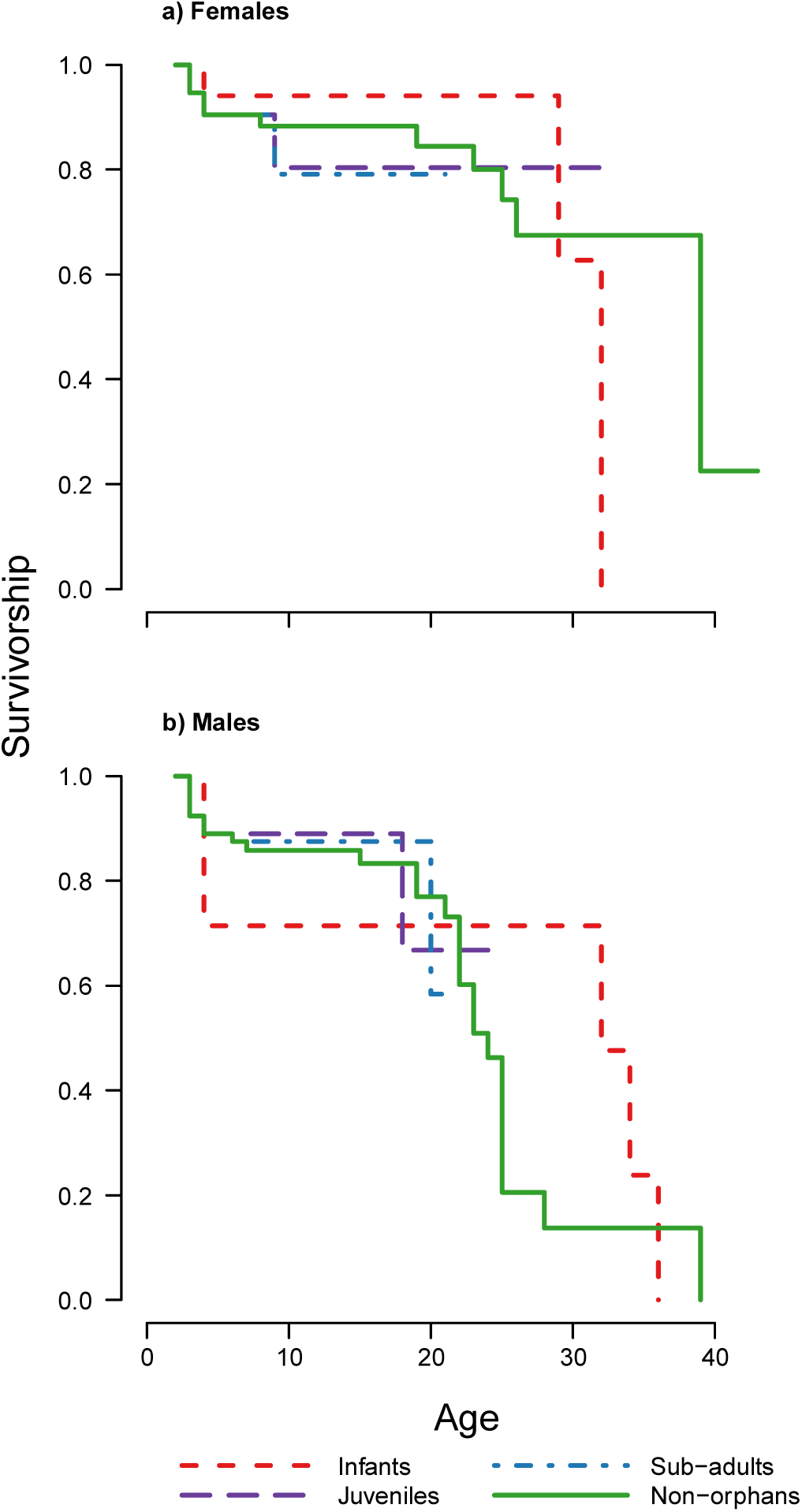
Survivorship curves for each of the four maternal loss categories for each sex. Plots show the proportion of surviving a) females and b) males, that suffered maternal loss as infants, juveniles and sub-adults compared to non-orphans that did not suffer maternal loss under the age of 8 years.

Bayesian survival trajectory analysis showed similar results, whereby the model with highest support for both sexes was the null model without orphan classes as covariates (Table 2).

**Table 2.**
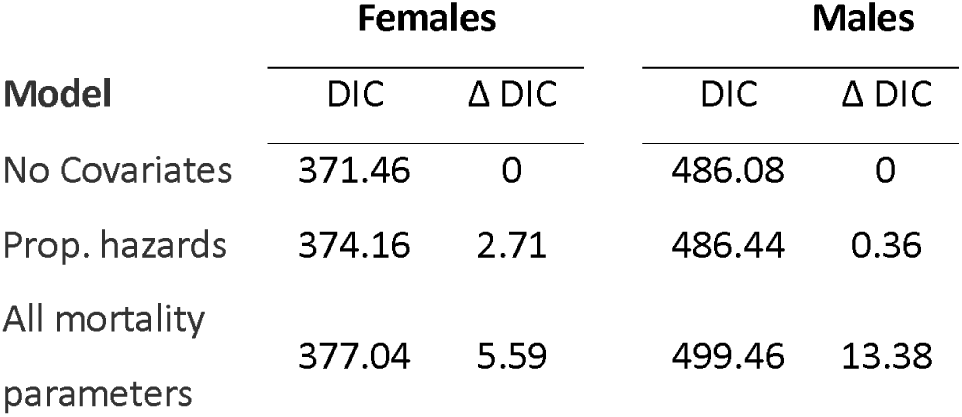
Deviance information criterion (DIC) for the three models tested. The Delta DIC shows the difference in DIC from the model with lowest DIC.

### Effect of maternal loss on dispersal

Female orphans were not significantly more likely to disperse from their natal group prior to first birth than female non-orphans (Table 3). However, there was a close to significant increase in dispersal for females that lost their mothers as juveniles or subadults. 37.5% of non-orphan females (n= 32) dispersed prior to their first birth (mean dispersal age ± SD: 7.96 ± 1.55) compared to 54.5% (n= 11) of infant orphans (dispersal age: 7.75 ± 0.60) and 75.0% (n= 8) of juvenile and subadult orphans (dispersal age: 8.21 ± 2.36). Only three males that reached the age of 16 had lost their mothers as infants, but all three remained in their natal group. Juvenile and subadult orphan males were significantly more likely to disperse before reaching 16 years (84.6%, n=13) than non-orphan males (37.5%, n=40, Table 3).

**Table 3.** The influence of age at maternal loss (infant or juvenile/subadult (J/SA)) relative to non-orphans on a female’s decision to disperse from their natal group prior to their first birth and a male’s decision to disperse prior to the age of 16, modelled using binomial generalized linear models.

|  | Females |  |  | Males |  |  |
| --- | --- | --- | --- | --- | --- | --- |
| | Est ± SE | t | P | Est ± SE | t | P |
| **Intercept** | -0.511 ± 0.365 | -1.399 | 0.162 | -0.511 ± 0.327 | -1.564 | 0.118 |
| **Infant** | 0.693 ± 0.707 | 0.980 | 0.327 | - | - | - |
| **J/SA** | 1.609 ± 0.894 | 1.799 | 0.072 | 2.216 ± 0.835 | 2.653 | **0.008** |

### Effect of maternal loss on female reproduction

Maternal loss had no significant effect on the age at which female orphans first gave birth (Supp. Table 2). The mean age at first birth (± SD) for non-orphans was 10.24 ± 1.61 compared to 9.72 ± 0.73 for those orphaned as infants and 9.67 ± 1.82 for those orphaned as juveniles or subadults. Accounting for age at first birth and dispersal, there was no evidence that maternal loss influenced whether a female’s first offspring survived infancy (Supp. Table 3). 51.5% of non-orphan females’ first-born offspring (n= 33) survived infancy compared to 60% (n= 10) of first-born offspring of those orphaned as infants and 57.1% (n=7) of first-born offspring of those orphaned as juveniles or subadults.

### Effect of maternal loss on male dominance attainment

As the oldest that a male first reached dominance was 22.99, we compared the proportion of males over the age of 23 that had attained dominance in each orphan class. 52.38% of non-orphan males had become the dominant male of a group for at least 6 months (n=21) compared to all three infant-orphaned males and no juvenile- or subadult-orphaned males (n=5). However, 5 out of the 7 juvenile- or subadult-orphaned males that had not yet reached 23 by the end of the study period or had died before reaching 23 years, had already become dominant.

### Social buffering of maternal loss

Both affiliative contact (t = 2.505, p =0.010) and proximity (t = 9.402, p <0.001) between orphans and other group members increased in the 6 months following maternal loss compared to the 6 months prior to maternal loss. Orphans’ weighted degree in the 2m proximity social network (the sum of the proportion of time spent within 2m of another gorilla) increased after maternal loss (t = 2.298, p = 0.029), despite losing their mother – commonly their closest social partner (Table 4). However, orphans did suffer a reduction in affiliative contact after maternal loss, with a decrease in their weighted degree in the contact network (t = -3.968, p <0.001).

**Table 4.** The percentage of gorilla orphans for which their mother was their closest social partner prior to maternal loss based on affiliative contact and proximity within 2 metres.

|  | Contact | Proximity |
| --- | --- | --- |
| Infants (n=9) | 88.8% | 77.8% |
| Juveniles (n=14) | 85.7% | 78.6% |
| Sub-adults (n=8) | 25.0% | 75.0% |

### Relationship changes following maternal loss

Affiliative contact and proximity between group members and orphans increased to a greater extent than between group members and other immature gorillas within the same group during the same time period (contact: t=6.660, p<0.001; proximity: t=2.743, p=0.006) (Figure 2, Supp. Table 4). The extent to which an orphan’s affiliative contact with and proximity to other group members increased following maternal loss did not differ depending on the orphan’s sex (Table 5). The increase in proximity with other group members following maternal loss was smaller for older orphans but this difference was not significant for affiliative contact (Table 5). This suggests that social support after maternal loss through proximity with other group members is lower for older orphans. Age-mates (those within 2 years age of the orphan) showed a greater increase in proximity after maternal loss relative to other group members. However, this was not the case for affiliative contact (Table 5). The change in relationship strength between maternal siblings after maternal loss depended on the age-sex class of the sibling (Supp. Figure 2). Subordinate adult males and subadult females had more affiliative contact with younger siblings following maternal loss but siblings in all other age-sex classes did not (Table 5). Both forms of social support (affiliative contact and proximity) showed the greatest increase from dominant males (Figure 2, Table 5). For affiliative contact this was significantly greater than all other age-sex classes, but for proximity the increase was only significantly greater than that of subordinate adult males.

**Table 5.**
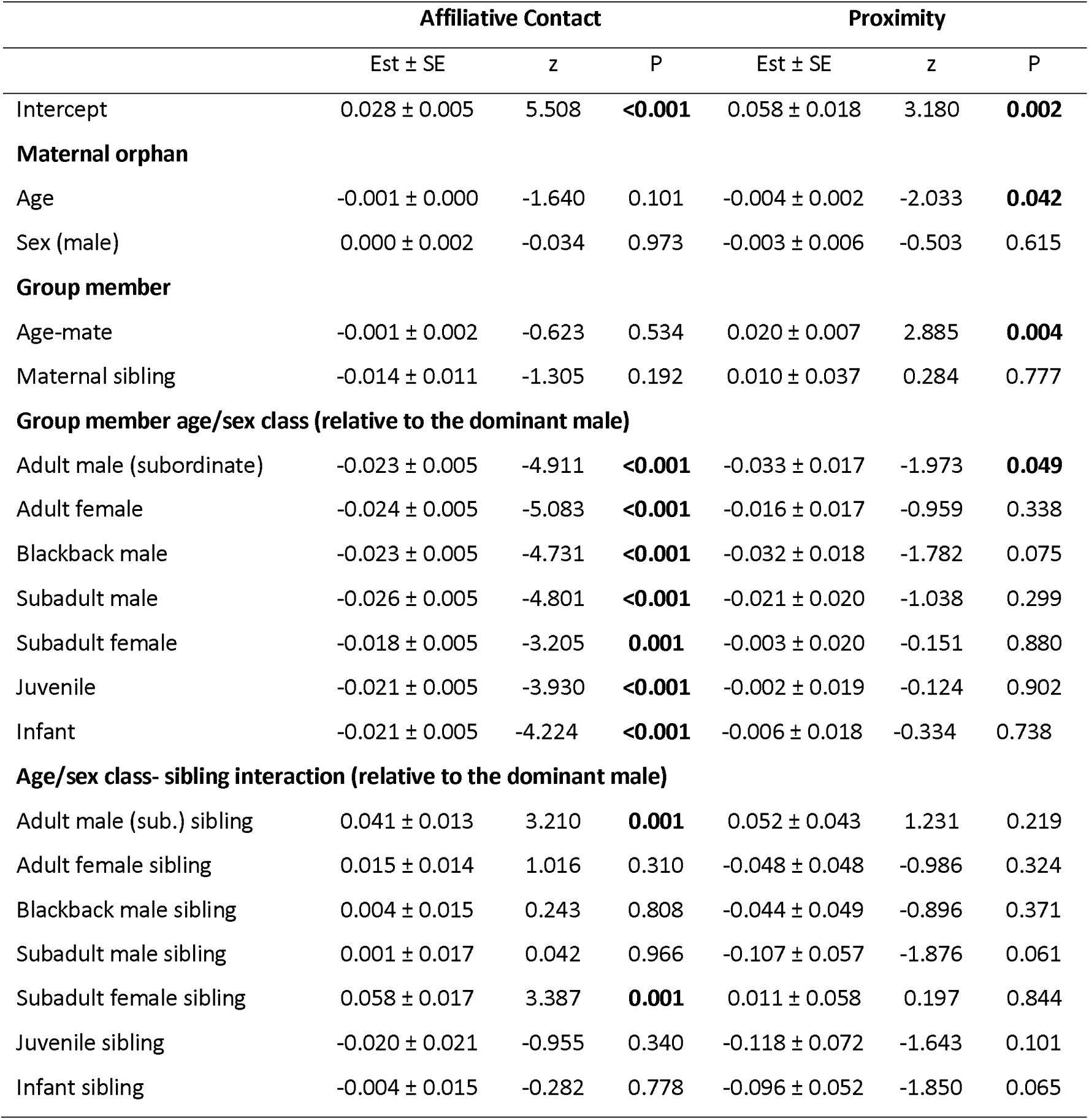
GAMM models predicting the change in dyadic relationship strength (SRI values for affiliative contact and proximity) between orphans and other group members following maternal loss

**Figure 2.**
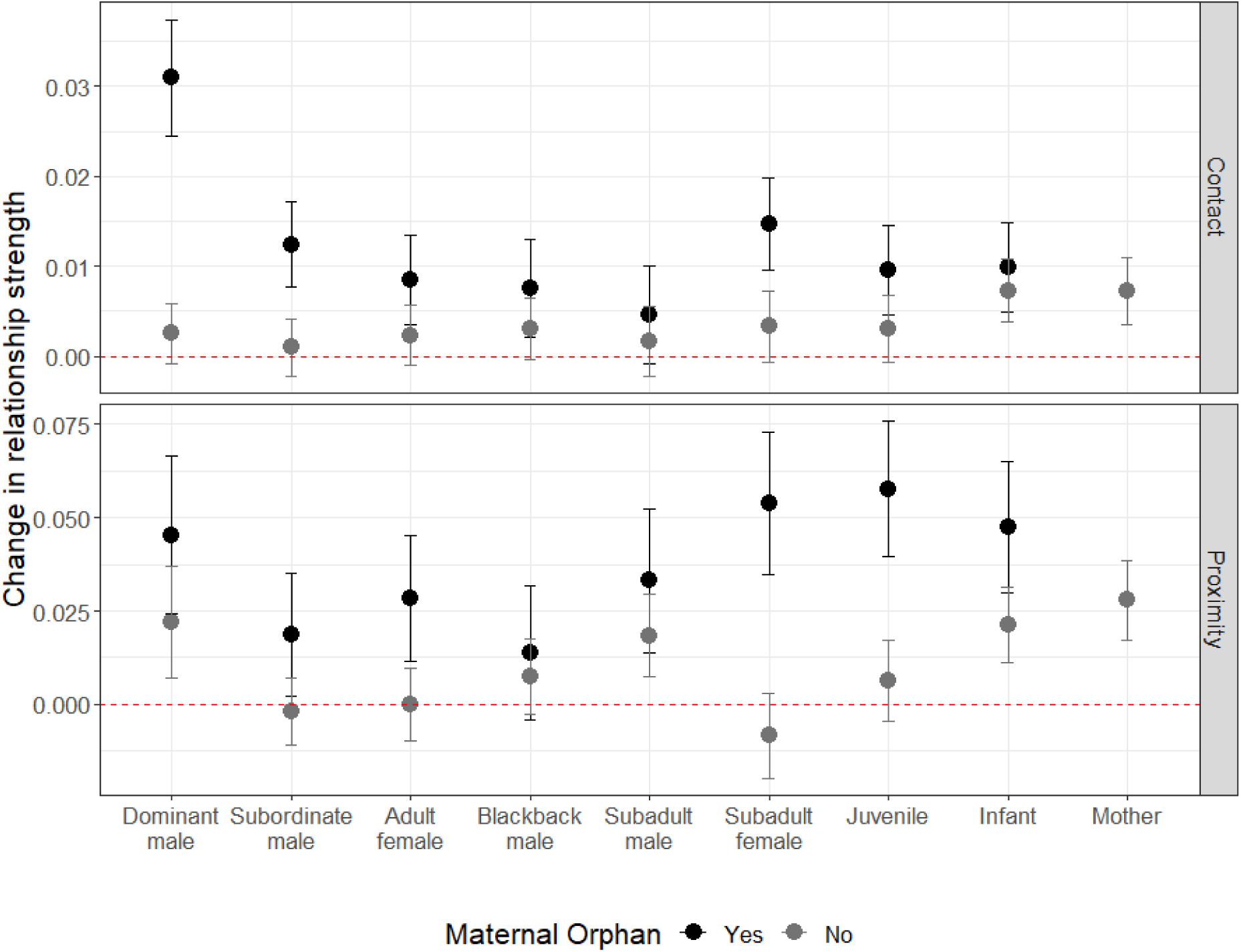
The change in relationship strength (SRI) between immature gorillas and other group members in both affiliative contact and proximity, between the 6 months prior to an incident of maternal loss and the 6 months post-maternal loss. Black points show values for orphans, grey points show values for immature gorillas within the same group that did not suffer maternal loss. Error bars indicate the standard error. Dashed red line indicates no change in relationship strength.

### Changes in the relationship with adult males following maternal loss

For 66.7% of orphans of known paternity (n=18), the dominant male at the time of maternal loss was their genetic father. Paternity did not influence the social support provided by adult males after maternal loss (contact: z=1.130, p=0.262; proximity: z=-0.552, p=0.583) but adult male maternal siblings increased both their affiliative contact and proximity more than non-siblings (contact: z=3.807, p<0.001; proximity: z=2.237, p=0.028). However, the increased social support from adult male maternal siblings relative to non-siblings was largely driven by an effect in subordinate males, whilst social support from dominant males did not differ greatly by kin relationship (Supp. Table 5, Supp. Fig. 3).

## Discussion

In contrast to many social mammals with extended periods of mother-offspring co-residence^7,8,10^, in mountain gorillas, we found no evidence for higher mortality in immature offspring of either sex following maternal loss. Whilst our sample size of 59 orphans between the ages of 2 and 8 years may limit our ability to detect relatively weak effects on survival, significant effects have been found in other species with comparable sample sizes^8^. This suggests that either maternal loss after offspring have reached 2 years does not reduce survival in mountain gorillas, or that the reduction in survival is considerably weaker than that observed in other species and therefore undetectable within our sample.

The effect of maternal loss on dispersal decisions differed depending on the age at which maternal loss occurred. The three males orphaned as infants remained in their natal group and females orphaned as infants were not significantly more likely to disperse than non-orphans. In contrast, both males and females orphaned as juveniles or subadults were more likely to disperse although for females this effect was not quite significant. This suggests that there may be some benefits to co-residence with mothers for both sexes that don’t directly translate to survival, but like the faster maturation and higher dominance in female chimpanzees^27,28^ or greater mating opportunities in male bonobos^16^, could influence lifetime reproductive success. Alternatively, potential benefits may not be specific to the mother-offspring bond. Instead, the loss of a key social partner, close to dispersal age may reduce the benefits of remaining in a group. This might better explain the age-dependent influence of maternal loss, as by the time those suffering maternal loss as infants approach an age at which dispersal could occur, they may have compensated for the loss of such a key social partner through strengthening their relationships with other group members. Our analyses also suggest that the strengthening of relationships with group members post-maternal loss may be greater for younger individuals, which could further reduce the likelihood that they disperse later in life.

We found no significant effect of maternal loss on the elements of female reproduction we investigated. Female orphans gave birth slightly younger, but despite this, their first offspring was marginally more likely to survive infancy. Whilst neither effect was significant, the direction of the effect, with earlier first birth, is consistent with the stress acceleration hypothesis: that those suffering from early caregiving adversity may show accelerated development as an adaptive strategy to low parental care^61,62^. Female dispersal has also not been found to cause any reproductive delays in mountain gorillas, suggesting that the potentially higher likelihood for those suffering maternal loss as juveniles or subadults to disperse is unlikely to reduce their reproductive capacity^63^.

Male reproduction was harder to assess due to limited paternity data and later sexual maturity. However, as dominant males usually sire the majority of offspring, even within multi-male groups^53–55^, analyses of male dominance status provided some insight into the reproductive success of male orphans. Although only around half of males ever reached dominant male status, some males in both orphan age categories were able to do so, demonstrating that this is possible in the absence of maternal support. Small sample sizes limited the extent of analysis possible, but maternal loss appeared to have differing effects depending on the age of maternal loss, with infant-orphaned males more likely and juvenile- or subadult-orphaned males less likely to become the dominant male of a group. This mirrors the effect of maternal loss on dispersal decisions. Given that dispersing males are less likely to become dominant^47^ and that modelling suggests dispersing males suffer a 50% reduction in lifetime reproductive success^64^, it is likely that males suffering maternal loss as juveniles or subadults may suffer from reduced siring opportunities throughout their lives.

Overall, these findings suggest there are no strong negative fitness consequences for maternal loss in female mountain gorillas over 2 years of age, although we cannot rule out longer-term effects on lifetime reproduction. In males, maternal loss after 2 years does not appear to influence survival but maternal loss as juveniles or subadults (4-8 years) could lower their future reproductive success. This lack of effect on survival reflects patterns found in human populations with natural fertility and mortality, where the negative effect of maternal loss on survival declines substantially with age and in many cases disappears entirely over the age of two^37^. However, it is contrary to research in most other social mammals with long periods of mother-offspring co-residency^1,5,7–9,15,18,32,36^. In these species, one of the major mechanisms posited to link maternal loss with increased mortality is through the social adversity caused by mother absence. Specifically, mother absence is found to reduce social integration^5,21^, competitivity^16,22^, dominance rank^6,18–20^ and opportunities for social learning^8,23,24,26^. But the lack of increased mortality in gorillas and certain human populations suggests that mother absence does not always lead to social adversity in mammals with extended maternal care.

When immature mountain gorillas suffered maternal loss, we found that their social relationships with other group members were strengthened, increasing their social integration within the group. The increase in proximity-based relationship strength with other group members was actually great enough to outweigh the loss of their mother, although this was not the case for relationships based on contact. In contrast to fission-fusion social systems such as in chimpanzees, where group composition regularly changes, mountain gorilla orphans remain in stable cohesive social groups in which their social integration is not dependent on their mother – as shown by the strengthening of relationships following maternal loss. Their increased social integration within the larger cohesive group following maternal loss suggests that mountain gorilla orphans are unlikely to have fewer opportunities for social learning as they are regularly in close proximity to multiple group members. Gorillas also do not appear to require the complex feeding techniques such as nut-cracking or termite-fishing, for which close contact with mothers may be most beneficial^25,26^. Instead, common group-membership may be sufficient for learning the complex foraging behaviours observed in gorillas as groups travel and feed as a cohesive unit.

The greatest increases in relationship strength following maternal loss were found with the dominant male of the group, regardless of whether he was the genetic father. This suggests that unlike in elephants where maternal loss leads to weaker relationships with dominant group members, gorillas that suffer maternal loss instead have increased access to the most dominant member of their group. Whilst the cohesion of gorilla social groups already limits the extent to which orphans are likely to suffer from reduced access to resources or knowledge, the strengthening of social relationships, particularly with the dominant male is likely to further buffer any potential reduction to their competitivity or future dominance rank within the group. Other cohesive social groups in which the effects of maternal loss have been studied have primarily been matriarchal, with groups led by older females e.g. elephants^32,36^ or where strong female dominance hierarchies are inherited e.g. spotted hyenas^9,18^ and cercopithecine species such as the savannah baboon^21^. In contrast, males are the dominant sex in gorillas^65^. Female dominance hierarchies are relatively weak, and the common dispersal of both sexes means maternal support is unlikely to provide considerable benefits for dominance to offspring of either sex. Orphaned gorillas are therefore unlikely to suffer costs in this regard. However, for males that remain in their natal group it is possible that stronger relationships with the dominant male, such as those developed post-maternal loss could aid dominance acquisition. For males that lose their mothers at particularly early ages it is possible this could ultimately improve their chances of inheriting dominance of their natal group^47,64^. Although, due to the rarity of such events (3 cases in 53 years) there may never be a large enough sample size to thoroughly investigate such a hypothesis.

The lack of paternity discrimination in the support provided by adult males after maternal loss is consistent with previous findings on paternal care in mountain gorillas, where the highest ranking males sire the majority of offspring and provide the most care, regardless of paternity^59,60^. As observed in chimpanzees^33^, support from some maternal siblings also appears to occur, with relationships between subordinate adult males and subadult females, and their younger siblings strengthening following maternal loss. In these two classes, caring for siblings may have additional benefits to those of inclusive fitness^66^. Care of infants by adult male mountain gorillas has been found to be linked with dramatically higher reproductive success. Males in the top tertile for showing affiliative behaviour towards infants, were found to sire 5.5 times more infants than those in the bottom tertile, even after accounting for rank^59^. This suggests that females may prefer males that demonstrate more caring behaviour towards infants, and that subordinate male kin that increase their care of maternal siblings following maternal loss may additionally increase their own reproductive success. Young, nulliparous female siblings, particularly subadult females may also benefit through developing parental experience that may improve their own reproductive success^66^.

In many human populations, care from other family members is believed to buffer the negative consequences of maternal loss, but the identity of these carers can vary greatly between populations. In rural Gambia, elder sisters and maternal grandmothers increased offspring survival but not fathers, other grandparents or brothers^67^. In contrast, in the Ache of Paraguay the loss of fathers significantly impacted offspring survival but the loss of grandparents or adult siblings did not^41^. Support from family members in rearing offspring is thought to be a human universal but the composition of those families and the specific family members involved in cooperative care appears to be flexible and responsive to ecological conditions^37^. In killer whales, as observed in many human populations, grandmothers have been found to influence offspring survival and this additional support could in part contribute to the lack of reduced survival for younger female killer whales after maternal loss^10,42^. In mountain gorillas, grandparents and their grand-offspring are rarely in the same social group as females often transfer between groups multiple times within their lives^46^. Instead, fathers (or those with a high likelihood of paternity), siblings and age-mates seem to play a key role in buffering the detrimental effects of maternal loss. What appears to set humans, gorillas and potentially killer whales apart from other species with extended maternal care where high fitness costs to maternal loss are observed, is the potential for cooperative care from within the social group.

## Conclusion

We found that immature mountain gorillas do not appear to face increased social adversity or a detectable reduction in fitness following maternal loss. It is not yet possible to demonstrate the direct link between the strengthening of relationships with other group members after maternal loss and the absence of fitness costs to maternal loss. However, our analyses show that at least in the short term, a key mechanism by which maternal loss is hypothesized to lead to reduced survival and fitness in other social species – social adversity - does not apply in mountain gorillas over the age of 2 years. The social support provided by other group members within mountain gorillas’ cohesive social groups, particularly from dominant males, age-mates and siblings, appears to buffer against the negative consequences of maternal loss.

In mountain gorillas, like humans^37–40,43–45^ social support appears to come from a number of group or family members. This could provide a buffer to the loss of any single relationship, even one as important as the mother-offspring relationship, once an individual is nutritionally independent. In the absence of nepotistic matriarchal dominance hierarchies and when social buffering is possible due to cohesive strongly bonded social groups, it may matter less who is providing care as long as care is provided.

## Methods

### Demographic data

Mountain gorillas in the Volcanoes National Park, Rwanda, have been monitored almost continuously by the Dian Fossey Gorilla Fund’s Karisoke Research Center since 1967. Habituated mountain gorilla groups are monitored daily by field teams who collect data on demography, behaviour, ranging and health. From 1967 to 2015 (inclusive), 59 immature mountain gorillas (28 males, 31 females) suffered maternal loss between the ages of 2 and 8 through the death or permanent transfer of their mother.

Gorillas were classified as infants up to 4 years of age, as juveniles from 4 to 6 years of age and as subadults from 6 to 8 years of age^57^. From 8 years of age females were classified as adults. Males were classified as blackbacks from 8 to 12 years. From 12 years, males were classified as either subordinate adult males or dominant adult males from their dominance hierarchy. Male dominance hierarchies were based on displacements and avoidances using the Elo-rating method^68,69^ (Supp. Fig. 1). Dominance hierarchies were calculated using the R package EloRating, version 0.43^70^ as described by Wright *et al*.^65^. Only one adult male was classified as dominant in each group at a given time. Dominant males were those with the highest dominance status unless they were the only adult male in the group in which case they were automatically classified as dominant.

### Survival

The youngest infant to survive maternal loss was a 2.45-year-old female. Our dataset included only 1 infant younger than this that suffered maternal loss at 0.67 years and died after 1 day. Three infants aged 1.91, 2.42 and 2.52 became separated from their mothers after a suspected poaching encounter. During this separation they travelled with a small number of group members not including their mothers. The 1.91-year-old died after 6 days of separation. The 2.42-year old died after 9 days of separation. The 2.52-year-old survived until they were reunited with their mother and the larger group 18 days after the initial separation. These infants were not included in the datasets as their mothers did not permanently transfer or die but combined with the maternal loss dataset, they suggest that infant mountain gorillas cannot survive independently from their mothers under the age of at least 2. We therefore investigated the effect of maternal loss after the age of 2.

To determine the effect of maternal loss on survival, we carried out a Cox-proportional hazards analysis separating individuals over the age of 2, based on four general age classes of maternal loss: a) infants (n=23), b) juveniles (n=23), c) sub-adults (n=13) and d) non-orphans (n=90), where mothers did not die or leave the group before the individual reached 8 years of age. First, we ran a Cox-proportional hazards model for each sex with a time-varying covariate for the age at maternal loss and using the four classes as covariates. Due to the small sample size for the sub-adult class, we merged this with the juvenile class into a single juvenile/sub-adult class and ran a new set of Cox-proportional hazards models on these new classes. In all cases we truncated the analysis to start at age 2.

In order to verify our results and to account for the uncertainty in some of the dates of birth, we ran a Bayesian survival trajectory analysis^71,72^ for each sex truncated at the age of maternal loss for all orphans, and at 2 years of age for non-orphans. We used orphan status as a binary covariate (orphan vs non-orphan) and, using the Siler mortality model^73^ for the baseline mortality, we tested three models: a) no covariates (i.e. null model where all individuals have the same hazard rate); b) proportional hazards (where mortality differs proportionally between orphan classes); c) covariates modifying all Siler mortality parameters (where each orphan class has a different age-specific mortality). We used deviance information criterion for model fit and selection^74,75^. Model b) is equivalent to the Cox-proportional hazards model. However, these tests facilitate further exploration of the hypotheses on the effect of maternal loss on mortality, namely that there is no effect (model a) or that the entire age-specific trajectory of mortality changes for each category.

### Female dispersal, age of first birth and survival of first offspring

Between 1967 and 2019, 66 females gave birth to what was known to be their first offspring. For 53 of these females, their age could be accurately estimated within a 90-day period. For these individuals, we extracted their age at first birth, whether they dispersed from their natal group prior to first birth and whether they had suffered maternal loss when immature, from the long-term database. Maternal loss was investigated with the age classes described above with juvenile and subadult classes merged due to small sample sizes. We investigated the effect of maternal loss on the decision of females to disperse from their natal group prior to their first birth using a binomial generalized linear model. We investigated the effect of both maternal loss and dispersal prior to first birth on age at first birth using a generalized linear model with a gaussian distribution. Due to the positive skew of age at first birth, we used the square root of age at first birth minus 8 (the earliest recorded age at first birth) as the response variable. We checked Q-Q plots to verify the normal distribution of residuals. Finally, we examined survival of each female’s first offspring through infancy (1: survived to 4 years, 0: died before reaching 4 years) according to age at first birth, dispersal prior to first birth (1: yes, 0: no) and maternal loss, with merged juvenile and subadult classes as above, using a binomial generalized linear model. Multicollinearity of all models with multiple variables was checked using variance inflation factors.

### Male dispersal and dominance

Between 1967 and 2019, 56 males whose age could be accurately estimated within a 90-day period, reached the median age of dispersal (16 years). We used a binomial generalized linear model to predict whether each of these males dispersed from their natal group prior to this age according to maternal loss classes (as above). To investigate the effect of maternal loss on dominance, we gave males that became the dominant male of a stable group for at least 6 consecutive months a dominance score of 1. This included adult males of single-male groups and the most dominant male of multi-male groups based on Elo-ratings (Supp. Fig. 1). Males that never reached dominance or only transiently (for <6 months) received a score of 0. We recorded the age at which a male first became dominant for those for which this could be accurately estimated within a 90-day period. This represented the age at which they first successfully attracted and retained a female to join their group, the age at which they split from their natal group with at least one adult female to form a new group, or the age at which their Elo-rating surpassed that of all other adult males in their group, if those groups remained independent and did not disintegrate within 6 months of that date. The mean age (± standard deviation) at which a male reached dominance was 17.88 ± 2.56. The median was 17.29. The oldest age at which a male first became dominant was 22.99. Therefore, to investigate the influence of maternal loss on dominance status we analysed only males that had survived and remained in the study population until at least 23 years (n=40). We did not attempt to statistically examine the effects of maternal loss on dominance status due to small sample sizes.

### Social network analysis

Gorilla groups were monitored for up to 4 hours daily and all gorillas were individually identified by physical characteristics. Behavioural data were collected on each group member via 50-minute focal sampling, with scan sampling completed every 10 minutes to record all gorillas within 2 metres of the focal individual and all gorillas in physical contact with the focal individual. Contact represented prolonged affiliative contact e.g. resting or feeding in contact, grooming or playing, and excluded physical aggression.

The social response of group members to an incident of maternal loss was investigated in 31 of the 59 total cases – those that suffered maternal loss after 2003 for which adequate social behavior data was available (more than 12 focal scans of the individual were recorded in the 6 months prior to maternal loss and the 6 months after maternal loss). For each case of maternal loss, two types of weighted social network were constructed based on a) 2m proximity and b) affiliative contact. For each type, edge values of the networks were calculated using the Simple Ratio Index (SRI)^76^. These values represented the proportion of time two individuals were either within 2m of each other or in physical contact. For example, in the contact network a value of 1 would indicate that the two individuals were in physical contact every time they were observed, whilst 0 would indicate that they were never observed in physical contact. Both network types were constructed for two time periods for each case of maternal loss: pre-maternal loss (the 6 months leading up to maternal loss) and post-maternal loss (the 6 months immediately after maternal loss). Social networks were constructed using all focal scans during these time periods.

### Social buffering of maternal loss

We extracted SRI edge values (representing the strength of relationship between a pair of gorillas) between the orphan and all other group members. We also extracted SRI edge values between all immature gorillas (aged 2-8) within the same group that had not suffered maternal loss and all other group members, for the same time periods. Only edge values involving gorillas for which more than 12 focal scans were available in both of the 6-month periods (pre- and post-maternal loss) were analysed^77^. This meant that only the social relationships with group members that were present both pre- and post-maternal loss were analysed, except for the mother-offspring relationships pre-maternal loss which were extracted separately. The mean (±sd) number of focal scans used to estimate edge values pre-maternal loss was 144.81 ± 90.10 and post maternal loss was 149.16 ± 88.56.

We used paired t-tests to compare the relationship strength (from both contact and 2m proximity) between an orphan and each group member pre- and post-maternal loss (n=755). Due to the non-independence data points, p-values were calculated via permutations using the ‘broman’ R package with 10,000 permutations which permuted the direction of change between paired samples. We calculated the weighted degree of an orphan in each network type pre-maternal loss as the sum of their SRI edge values with all other group members and the SRI value of the mother-offspring bond during the pre-maternal loss period. We calculated the weighted degree post-maternal loss as the sum of SRI edge values with all other group members (without the mother) during the post-maternal loss period. We used paired t-tests to compare the weighted degree of an immature gorilla pre- and post-maternal loss in both their contact and 2m proximity networks (n=31). P-values were calculated using permutations as above.

### Relationship changes following maternal loss

We estimated the change in relationship strength between an immature gorilla (both orphans and non-orphans) with other group members following an incident of maternal loss within the group as the change in SRI value between periods (SRI post-maternal loss – SRI pre-maternal loss) in both network types (contact and proximity). Generalized additive mixed models (GAMMs) in the ‘gamm4’ R package were used to predict this change for both affiliative contact and proximity which enabled the non-independence of network data to be accounted for through random effect structures and sampling variation to be accounted for using a smoothing term. We ran an initial set of GAMMs on the change in SRI values for relationships involving the orphans or other immature gorillas within the same group that had not suffered maternal loss, excluding mother-offspring relationships. This was to verify that changes in the relationships of orphans with other group members were significantly different to those observed for other immature gorillas present within the same group during that period. These GAMMs included the identity of the immature gorilla as a random factor, nested within the specific maternal loss incident (group and time period), nested within the group. The identity of the other group member was included as a further random factor. The mean number of focal scans used to estimate the SRI value across both time periods was included as a smoothing term in the model to account for any biases from sampling intensity. The age and sex of the immature gorilla were included as fixed factors, along with the age-sex class of the other group member, whether the immature gorilla suffered maternal loss during this period (1:Yes, 0: No), and the interaction between maternal loss and the age-sex class of the other group member. Non-orphan relationships where the other group member was an orphan were excluded from the analysis.

We then ran GAMMs on only the relationships involving orphans to investigate in more detail how these changed following maternal loss. The identity of the orphan was included as a random factor, nested within the group. The identity of the other group member was included as a further random factor. The mean number of focal scans used to estimate the SRI value across both time periods was again included as a smoothing term. The age and sex of the orphan were included as fixed factors to predict the social response of group members, as well as whether those group members were maternal siblings (1:Yes, 0:No) and whether those group members were age-mates (1: <2 years age difference with the orphan, 0: ≥2 years age difference). This 2-year cut-off was chosen to be consistent with the width of age categories, such that age-mates would be within the same age class during large proportions of their immature life. The age-sex class of the other group member at the date of maternal loss and the interaction between this and maternal sibling status were also included as fixed factors for predicting the change in relationship.

### Adult male relationship changes following maternal loss

Paternity was known for 18 of the 31 orphans for which social data was available, from a previous study^78^. To investigate the influence of paternity, we ran an additional set of GAMMS to predict the change in relationship (both contact and proximity) between adult males and orphans following maternal loss (n=121). The identity of the orphan and the identity of the adult male were included as random factors. The mean number of focal scans was included as a smoothing term. Orphan age, orphan sex, male dominance, and the binary kinship variables: maternal sibling and paternity, were included as fixed factors. The interaction between dominance and sibling status was also included in the model, but the interaction between dominance and paternity could not be, due to possible issues of multicollinearity (variance inflation factors >3).

## Acknowledgements

We are grateful to the Rwandan Development Board (RDB) for their long-term support of the Karisoke Research Center and thank the Karisoke field staff for collection of the data. We are thankful to Edward Wright for his support with the Elo-rating analysis for male dominance hierarchies and grateful to Lauren Brent and her research group for providing insightful feedback and discussion on the project. We thank Elizabeth Lonsdorf for her valuable feedback on the manuscript.

## Author contributions

WE conceived the project, which was further developed by all authors. Data collection was managed by WE, VV and TSS. Data was compiled by WE and VV and verified by REM. Analysis was done by REM, FC and WE. REM led the writing with contributions from FC, WE and TSS and final approval from all authors.

## References

1. Snyder-Mackler, N. et al. Social determinants of health and survival in humans and other animals. Science (80-.). 368, eaax9553 (2020).

2. Mitani, J. C., Call, J., Kappeler, P. M., Palombit, R. A. & Silk, J. B. The Evolution of Primate Societies. The Evolution of Primate Societies (2013). doi: 10.7208/chicago/9780226531731.001.0001

3. Nowak, R., Porter, R. H., Lévy, F., Orgeur, P. & Schaal, B. Role of mother-young interactions in the survival of offspring in domestic mammals. Reviews of Reproduction (2000). doi: 10.1530/ror.0.0050153

4. Clutton-Brock, T. H. The Evolution of Parental Care. (Princeton University Press, 1991).

5. Tung, J., Archie, E. A., Altmann, J. & Alberts, S. C. Cumulative early life adversity predicts longevity in wild baboons. Nat. Commun. (2016). doi: 10.1038/ncomms11181

6. Strauss, E. D., Shizuka, D. & Holekamp, K. E. Juvenile rank acquisition is associated with fitness independent of adult rank. Proc. R. Soc. B Biol. Sci. (2020). doi: 10.1098/rspb.2019.2969

7. Andres, D. et al. Sex differences in the consequences of maternal loss in a long-lived mammal, the red deer (Cervus elaphus). Behav. Ecol. Sociobiol. (2013). doi: 10.1007/s00265-013-1552-3

8. Stanton, M. A., Lonsdorf, E. V., Murray, C. M. & Pusey, A. E. Consequences of maternal loss before and after weaning in male and female wild chimpanzees. Behav. Ecol. Sociobiol. (2020). doi: 10.1007/s00265-020-2804-7

9. Watts, H. E., Tanner, J. B., Lundrigan, B. L. & Holekamp, K. E. Post-weaning maternal effects and the evolution of female dominance in the spotted hyena. Proc. R. Soc. B Biol. Sci. (2009). doi: 10.1098/rspb.2009.0268

10. Foster, E. A. et al. Adaptive prolonged postreproductive life span in killer whales. Science (80-.). (2012). doi: 10.1126/science.1224198

11. Holt-Lunstad, J., Smith, T. B. & Layton, J. B. Social relationships and mortality risk: A meta-analytic review. PLoS Medicine (2010). doi: 10.1371/journal.pmed.1000316

12. Fairbanks, L. A. Maternal investment throughout the life span in Old World monkeys. in Old World Monkeys (2009). doi: 10.1017/cbo9780511542589.014

13. Altmann, J. & Alberts, S. C. Growth rates in a wild primate population: Ecological influences and maternal effects. Behav. Ecol. Sociobiol. (2005). doi: 10.1007/s00265-004-0870-x

14. Greenwood, P. J. Mating systems, philopatry and dispersal in birds and mammals. Anim. Behav. 28, 1140–1162 (1980).

15. Nakamura, M., Hayaki, H., Hosaka, K., Itoh, N. & Zamma, K. Brief Communication: Orphaned male Chimpanzees die young even after weaning. Am. J. Phys. Anthropol. (2014). doi: 10.1002/ajpa.22411

16. Surbeck, M. et al. Males with a mother living in their group have higher paternity success in bonobos but not chimpanzees. Current Biology (2019). doi: 10.1016/j.cub.2019.03.040

17. Surbeck, M., Mundry, R. & Hohmann, G. Mothers matter! Maternal support, dominance status and mating success in male bonobos (Pan paniscus). in Proceedings of the Royal Society B: Biological Sciences (2011). doi: 10.1098/rspb.2010.1572

18. East, M. L. et al. Maternal effects on offspring social status in spotted hyenas. Behav. Ecol. (2009). doi: 10.1093/beheco/arp020

19. Maestripieri, D. & Mateo, J. M. Maternal Effects in Mammals. (University of Chicago Press, 2009). doi: 10.7208/chicago/9780226501222.003.0014

20. Lea, A. J., Learn, N. H., Theus, M. J., Altmann, J. & Alberts, S. C. Complex sources of variance in female dominance rank in a nepotistic society. Anim. Behav. (2014). doi: 10.1016/j.anbehav.2014.05.019

21. Archie, E. A., Tung, J., Clark, M., Altmann, J. & Alberts, S. C. Social affiliation matters: Both same-sex and opposite-sex relationships predict survival in wild female baboons. Proc. R. Soc. B Biol. Sci. (2014). doi: 10.1098/rspb.2014.1261

22. Samuni, L. et al. Maternal effects on offspring growth indicate post-weaning juvenile dependence in chimpanzees (Pan troglodytes verus). Front. Zool. 17, 1–12 (2020).

23. Foley, C., Pettorelli, N. & Foley, L. Severe drought and calf survival in elephants. Biol. Lett. (2008). doi: 10.1098/rsbl.2008.0370

24. Brent, L. J. N. et al. Ecological knowledge, leadership, and the evolution of menopause in killer whales. Curr. Biol. (2015). doi: 10.1016/j.cub.2015.01.037

25. Lonsdorf, E. V., Eberly, L. E. & Pusey, A. E. Sex differences in learning in chimpanzees. Nature (2004). doi: 10.1038/428715a

26. Estienne, V., Cohen, H., Wittig, R. M. & Boesch, C. Maternal influence on the development of nut-cracking skills in the chimpanzees of the Taï forest, Côte d’Ivoire (Pan troglodytes verus). Am. J. Primatol. (2019). doi: 10.1002/ajp.23022

27. Walker, K. K., Walker, C. S., Goodall, J. & Pusey, A. E. Maturation is prolonged and variable in female chimpanzees. J. Hum. Evol. (2018). doi: 10.1016/j.jhevol.2017.10.010

28. Foerster, S. et al. Chimpanzee females queue but males compete for social status. Sci. Rep. (2016). doi: 10.1038/srep35404

29. Zipple, M. N., Archie, E. A., Tung, J., Altmann, J. & Alberts, S. C. Intergenerational effects of early adversity on survival in wild baboons. Elife (2019). doi: 10.7554/elife.47433

30. Hamilton, W. J., Busse, C. & Smith, K. S. Adoption of infant orphan chacma baboons. Anim. Behav. (1982). doi: 10.1016/S0003-3472(82)80233-9

31. Engh, A. L. et al. Behavioural and hormonal responses to predation in female chacma baboons (Papio hamadryas ursinus). Proc. R. Soc. B Biol. Sci. (2006). doi: 10.1098/rspb.2005.3378

32. Goldenberg, S. Z. & Wittemyer, G. Orphaned female elephant social bonds reflect lack of access to mature adults. Sci. Rep. (2017). doi: 10.1038/s41598-017-14712-2

33. Reddy, R. B. & Mitani, J. C. Social relationships and caregiving behavior between recently orphaned chimpanzee siblings. Primates (2019). doi: 10.1007/s10329-019-00732-1

34. Firth, J. A. et al. Wild birds respond to flockmate loss by increasing their social network associations to others. Proc. R. Soc. B Biol. Sci. (2017). doi: 10.1098/rspb.2017.0299

35. Hobaiter, C., Schel, A. M., Langergraber, K. & Zuberbühler, K. ‘Adoption’ by maternal siblings in wild chimpanzees. PLoS One (2014). doi: 10.1371/journal.pone.0103777

36. Goldenberg, S. Z. & Wittemyer, G. Orphaning and natal group dispersal are associated with social costs in female elephants. Anim. Behav. (2018). doi: 10.1016/j.anbehav.2018.07.002

37. Sear, R. & Mace, R. Who keeps children alive? A review of the effects of kin on child survival. Evolution and Human Behavior (2008). doi: 10.1016/j.evolhumbehav.2007.10.001

38. Lahdenperä, M., Lummaa, V., Helle, S., Tremblay, M. & Russell, A. F. Fitness benefits of prolonged post-reproductive lifespan in women. Nature (2004). doi: 10.1038/nature02367

39. Sear, R., Mace, R. & McGregor, I. A. Maternal grandmothers improve nutritional status and survival of children in rural Gambia. Proc. R. Soc. B Biol. Sci. (2000). doi: 10.1098/rspb.2000.1190

40. Hurtado, A. M. & Hill, K. R. Paternal effect on offspring survivorship among ache and hiwi hunter-gatherers: Implications for modeling pair-bond stability. in Father-Child Relations: Cultural and Biosocial Contexts (1992). doi: 10.4324/9780203792063

41. Hill, K. & Hurtado, A. M. Ache Life History: The Ecology and Demography of a Foraging People. (Routledge, 2017). doi: 10.2307/2137817

42. Nattrass, S. et al. Postreproductive killer whale grandmothers improve the survival of their grandoffspring. Proc. Natl. Acad. Sci. U. S. A. (2019). doi: 10.1073/pnas.1903844116

43. Andersson, T., Högberg, U. & Åkerman, S. Survival of orphans in 19th century Sweden - The importance of remarriages. Acta Paediatr. Int. J. Paediatr. (1996). doi: 10.1111/j.1651-2227.1996.tb14198.x

44. Campbell, C. & Lee, J. Z. When husbands and parents die: widowhood and orphanhood in late Imperial Liaoning, 1789-1909. When Dad Died Individ. Fam. Coping with Distress Past Soc. (2002).

45. Bentley, G. R. & Mace, R. Substitute parents: Biological and social perspectives on alloparenting across human societies. Substitute Parents: Biological and Social Perspectives on Alloparenting across Human Societies (2009).

46. Robbins, A. M., Stoinski, T., Fawcett, K. & Robbins, M. M. Leave or conceive: natal dispersal and philopatry of female mountain gorillas in the Virunga volcano region. Anim. Behav. 77, 831–838 (2009).

47. Stoinski, T. S. et al. Proximate factors influencing dispersal decisions in male mountain gorillas, Gorilla beringei beringei. Anim. Behav. (2009). doi: 10.1016/j.anbehav.2008.12.030

48. Morrison, R., Groenenberg, M., Breuer, T., Manguette, M. & Walsh, P. Hierarchical Social Modularity in Gorillas. Proc. R. Soc. B Biol. Sci. (2019). doi: https://doi.org/10.1098/rspb.2019.0681

49. Mirville, M. O. et al. Low familiarity and similar ‘group strength’ between opponents increase the intensity of intergroup interactions in mountain gorillas (Gorilla beringei beringei). Behav. Ecol. Sociobiol. 72, (2018).

50. Morrison, R. E., Eckardt, W., Stoinski, T. S. & Brent, L. J. N. Comparing measures of social complexity: larger mountain gorilla groups do not have a greater diversity of relationships. [In Prep]

51. Gray, M. et al. Censusing the mountain gorillas in the Virunga Volcanoes: Complete sweep method versus monitoring. Afr. J. Ecol. (2010). doi: 10.1111/j.1365-2028.2009.01142.x

52. Robbins, M. M. A Demographic Analysis of Male Life History and Social Structure of Mountain Goril. Behaviour (1995). doi: 10.1163/156853995x00261

53. Nsubuga, A. M., Robbins, M. M., Boesch, C. & Vigilant, L. Patterns of paternity and group fission in wild multimale mountain gorilla groups. Am. J. Phys. Anthropol. 135, 263–274 (2008).

54. Stoinski, T. S. et al. Patterns of male reproductive behaviour in multi-male groups of mountain gorillas: Examining theories of reproductive skew. Behaviour (2009). doi: 10.1163/156853909x419992

55. Bradley, B. J. et al. Mountain gorilla tug-of-war: silverbacks have limited control over reproduction in multimale groups. Proc. Natl. Acad. Sci. U. S. A. 102, 9418–9423 (2005).

56. Eckardt, W., Fawcett, K. & Fletcher, A. W. Weaned age variation in the Virunga mountain gorillas (Gorilla beringei beringei): influential factors. Behav. Ecol. Sociobiol. (2016). doi: 10.1007/s00265-016-2066-6

57. Breuer, T., Hockemba, M. B. N., Olejniczak, C., Parnell, R. J. & Stokes, E. J. Physical maturation, life-history classes and age estimates of free-ranging western gorillas - Insights from Mbeli Bai, Republic of Congo. Am. J. Primatol. 71, 106–119 (2009).

58. Harcourt, A. H. & Greenberg, J. Do gorilla females join males to avoid infanticide? A quantitative model. Anim. Behav. (2001). doi: 10.1006/anbe.2001.1835

59. Rosenbaum, S., Vigilant, L., Kuzawa, C. W. & Stoinski, T. S. Caring for infants is associated with increased reproductive success for male mountain gorillas. Sci. Rep. (2018). doi: 10.1038/s41598-018-33380-4

60. Rosenbaum, S., Hirwa, J. P., Silk, J. B., Vigilant, L. & Stoinski, T. S. Male rank, not paternity, predicts male-immature relationships inmountain gorillas, Gorilla beringei beringei. Anim. Behav. 104, 13–24 (2015).

61. Ellis, B. J. Timing of pubertal maturation in girls: An integrated life history approach. Psychological Bulletin (2004). doi: 10.1037/0033-2909.130.6.920

62. Callaghan, B. L. & Tottenham, N. The Stress Acceleration Hypothesis: Effects of early-life adversity on emotion circuits and behavior. Current Opinion in Behavioral Sciences (2016). doi: 10.1016/j.cobeha.2015.11.018

63. Robbins, A. M., Stoinski, T. S., Fawcett, K. A. & Robbins, M. M. Does dispersal cause reproductive delays in female mountain gorillas? Behaviour (2009). doi: 10.1163/156853909x426354

64. Robbins, A. M. & Robbins, M. M. Fitness consequences of dispersal decisions for male mountain gorillas (Gorilla beringei beringei). Behav. Ecol. Sociobiol. 58, 295–309 (2005).

65. Wright, E. et al. Male body size, dominance rank and strategic use of aggression in a group-living mammal. Anim. Behav. (2019). doi: 10.1016/j.anbehav.2019.03.011

66. Riedman, M. L. The Evolution of Alloparental Care and Adoption in Mammals and Birds. Q. Rev. Biol. (1982). doi: 10.1086/412936

67. Sear, R., Mace, R. & McGregor, I. A. The effects of kin on female fertility in rural Gambia. Evol. Hum. Behav. 24, 25–42 (2003).

68. Albers, P. C. H. & De Vries, H. Elo-rating as a tool in the sequential estimation of dominance strengths. Animal Behaviour (2001). doi: 10.1006/anbe.2000.1571

69. Neumann, C. et al. Assessing dominance hierarchies: Validation and advantages of progressive evaluation with Elo-rating. Animal Behaviour (2011). doi: 10.1016/j.anbehav.2011.07.016

70. Neumann, C. & Lars, K. EloRating: Animal Dominance Hierarchies by Elo Rating. R Packag. version 0.43. (2014).

71. Colchero, F. & Clark, J. S. Bayesian inference on age-specific survival for censored and truncated data. J. Anim. Ecol. (2012). doi: 10.1111/j.1365-2656.2011.01898.x

72. Colchero, F., Jones, O. R. & Rebke, M. BaSTA: An R package for Bayesian estimation of age-specific survival from incomplete mark-recapture/recovery data with covariates. Methods Ecol. Evol. (2012). doi: 10.1111/j.2041-210X.2012.00186.x

73. Siler, W. A Competing-Risk Model for Animal Mortality. Ecology (1979). doi: 10.2307/1936612

74. Spiegelhalter, D. J., Best, N. G., Carlin, B. P. & Van Der Linde, A. Bayesian measures of model complexity and fit. J. R. Stat. Soc. Ser. B Stat. Methodol. (2002). doi: 10.1111/1467-9868.00353

75. Celeux, G., Forbesy, F., Robertz, C. P. & Titteringtonx, D. M. Deviance information criteria for missing data models. Bayesian Anal. (2006). doi: 10.1214/06-BA122

76. Whitehead, H. Analyzing Animal Societies: Quantitative Methods for Vertebrate Social Analysis. (University of Chicago Press, 2008).

77. Farine, D. R. & Whitehead, H. Constructing, conducting and interpreting animal social network analysis. J. Anim. Ecol. 84, 1144–63 (2015).

78. Vigilant, L. et al. Reproductive competition and inbreeding avoidance in a primate species with habitual female dispersal. Behav. Ecol. Sociobiol. (2015). doi: 10.1007/s00265-015-1930-0

