## Supplementary material for "Social groups buffer maternal loss in mountain gorillas": Supp.

**Supp. Table 1**. Cox-proportional hazards models showing the effects of the three maternal loss classes on survival. All results are relative to the non-orphan class.

|  | **Females** | |  |  | **Males** | |
| --- | --- | --- | --- | --- | --- | --- |
| **Age-class** | Mean ± SE | P | |  | Mean ± SE | P |
| Infants | 0.626 ± 0.585 | 0.284 | |  | -0.351 ± 0.549 | 0.523 |
| Juveniles/subadults | 0.927 ± 1.140 | 0.415 | |  | 0.025 ± 1.050 | 0.981 |

**Supp. Table 2.** The influence of age of maternal loss (infant or juvenile/subadult - J/SA) relative to non-orphans and dispersal prior to their first birth on a female’s age at first birth, modelled using a generalized linear model with a gaussian distribution.

|  | **Est ± SE** | **t** | **P** |
| --- | --- | --- | --- |
| **Intercept** | 1.495 ± 0.121 | 12.573 | <0.001 |
| **Dispersal** | -0.245 ± 0.197 | -1.239 | 0.222 |
| **Infant** | -0.154 ± 0.270 | -0.569 | 0.572 |
| **J/SA** | -0.552 ± 0.554 | -0.996 | 0.325 |
| **Dispersal:Infant** | 0.149 ± 0.382 | 0.390 | 0.698 |
| **Dispersal: J/SA** | 0.501 ± 0.611 | 0.820 | 0.416 |

**Supp. Table 3.** The influence of age at birth, dispersal and age of maternal loss (I – infant, J/S –juvenile/sub-adult) on whether a female’s first offspring survived infancy, modelled using a binomial generalized linear model.

|  | **Est ± SE** | **Z** | **P** |
| --- | --- | --- | --- |
| **Intercept** | -1.402 ± 2.092 | -0.670 | 0.503 |
| **Age at birth** | 0.158 ± 0.198 | 0.799 | 0.424 |
| **Dispersal** | -0.264 ± 0.618 | -0.428 | 0.669 |
| **Maternal Loss I** | 0.397 ± 0.751 | 0.528 | 0.598 |
| **Maternal Loss J/S** | 0.375 ± 0.905 | 0.414 | 0.679 |

**Supp. Table 4.** GAMM models predicting the change in dyadic relationship strength (SRI values for affiliative contact and proximity) between all immature gorillas (orphans and non-orphans) and other group members following an incident of maternal loss within the group.

|  | **Affiliative Contact** | | | **Proximity** | | |
| --- | --- | --- | --- | --- | --- | --- |
|  | Est ± SE | z | P | Est ± SE | z | P |
| Intercept | 0.006± 0.003 | 2.002 | 0.045 | 0.023 ± 0.014 | 1.635 | 0.102 |
| **Immature** | | | | | | |
| Maternal orphan | 0.021 ± 0.003 | 6.660 | **<0.001** | 0.029 ± 0.011 | 2.743 | **0.006** |
| Age | -0.001 ± 0.000 | -3.544 | **<0.001** | -0.002 ± 0.001 | -3.864 | **<0.001** |
| Sex (male) | 0.001 ± 0.001 | 1.025 | 0.306 | 0.006 ± 0.002 | 2.808 | **0.005** |
| **Group member age/sex class (relative to the dominant male)** | | | | | | |
| Adult male (subordinate) | -0.003 ± 0.002 | -1.491 | 0.136 | -0.017 ± 0.008 | -2.006 | **0.045** |
| Adult female | -0.002 ± 0.002 | -0.732 | 0.464 | -0.008 ± 0.009 | -0.936 | 0.349 |
| Blackback male | -0.001 ± 0.002 | -0.493 | 0.622 | -0.004 ± 0.009 | -0.418 | 0.676 |
| Subadult male | -0.002 ± 0.003 | -0.868 | 0.385 | 0.009 ± 0.010 | 0.864 | 0.388 |
| Subadult female | -0.001 ± 0.003 | -0.529 | 0.597 | -0.015 ± 0.011 | -1.399 | 0.162 |
| Juvenile | -0.001 ± 0.003 | -0.411 | 0.681 | -0.004 ± 0.010 | -0.394 | 0.693 |
| Infant | 0.003 ± 0.002 | 1.228 | 0.219 | 0.013 ± 0.009 | 1.377 | 0.169 |
| **Orphan - age/sex class interaction (relative to the dominant male)** | | | | | | |
| Adult male (subordinate) | -0.015 ± 0.004 | -4.238 | **<0.001** | -0.014 ± 0.012 | -1.131 | 0.258 |
| Adult female | -0.021 ± 0.003 | -6.306 | **<0.001** | -0.018 ± 0.011 | -1.599 | 0.110 |
| Blackback male | -0.023 ± 0.004 | -6.358 | **<0.001** | -0.036 ± 0.012 | -2.972 | **0.003** |
| Subadult male | -0.025 ± 0.004 | -6.200 | **<0.001** | -0.033 ± 0.014 | -2.418 | **0.016** |
| Subadult female | -0.015 ± 0.004 | -3.812 | **<0.001** | 0.009 ± 0.014 | 0.681 | 0.496 |
| Juvenile | -0.021 ± 0.004 | -5.386 | **<0.001** | 0.002 ± 0.013 | 0.148 | 0.882 |
| Infant | -0.025 ± 0.004 | -7.032 | **<0.001** | -0.024 ± 0.012 | -2.030 | **0.042** |

**Supp. Table 5.** GAMM models predicting the change in dyadic relationship strength (SRI values for affiliative contact and proximity) between adult males and orphans following maternal loss.

|  | Affiliative Contact | | | Proximity | | |
| --- | --- | --- | --- | --- | --- | --- |
|  | Est ± SE | Z | P | Est ± SE | z | P |
| Intercept | 0.008 ± 0.009 | 0.881 | 0.381 | 0.017 ± 0.029 | 0.604 | 0.548 |
| **Maternal orphan** | | | | | | |
| Age | -0.002 ± 0.001 | -1.051 | 0.296 | -0.003 ± 0.005 | -0.643 | 0.522 |
| Sex M | 0.002 ± 0.004 | 0.550 | 0.584 | 0.001 ± 0.013 | 0.048 | 0.962 |
| **Adult Male** | | | | | | |
| Father | 0.009 ± 0.008 | 1.130 | 0.262 | -0.014 ± 0.026 | -0.552 | 0.583 |
| Maternal sibling | 0.027 ± 0.007 | 3.807 | **<0.001** | 0.051 ± 0.023 | 2.237 | **0.028** |
| Dominant male | 0.008 ± 0.009 | 0.859 | 0.393 | 0.084± 0.030 | 2.768 | **0.007** |
| **Dominance-kin interaction** |  |  |  |  |  |  |
| Dominant sibling | -0.039 ± 0.013 | -2.932 | **0.004** | -0.074 ± 0.041 | -1.765 | 0.082 |

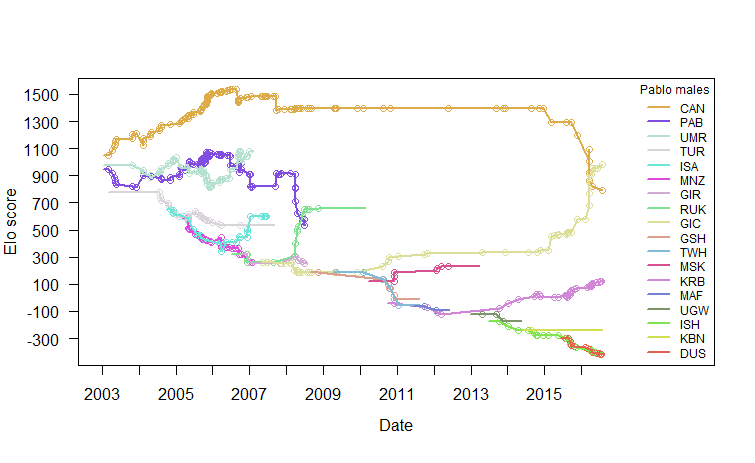

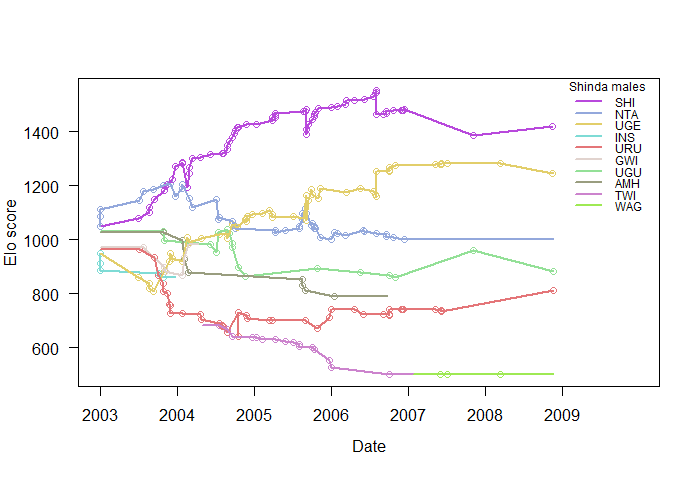

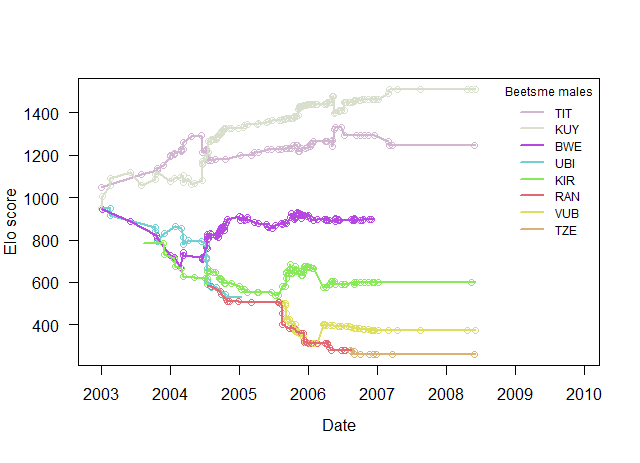

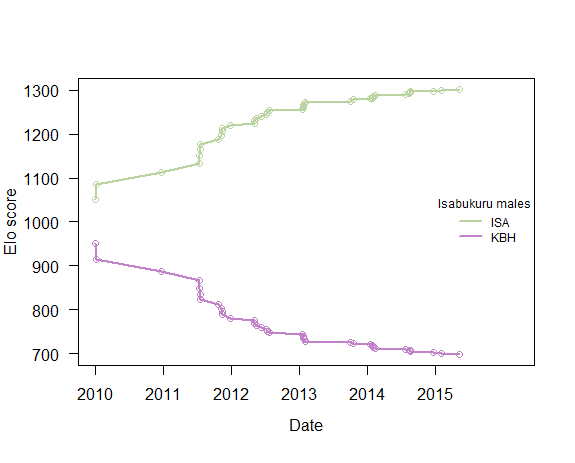

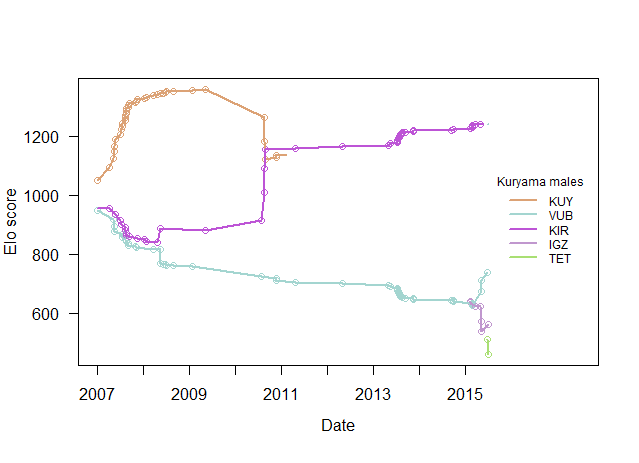

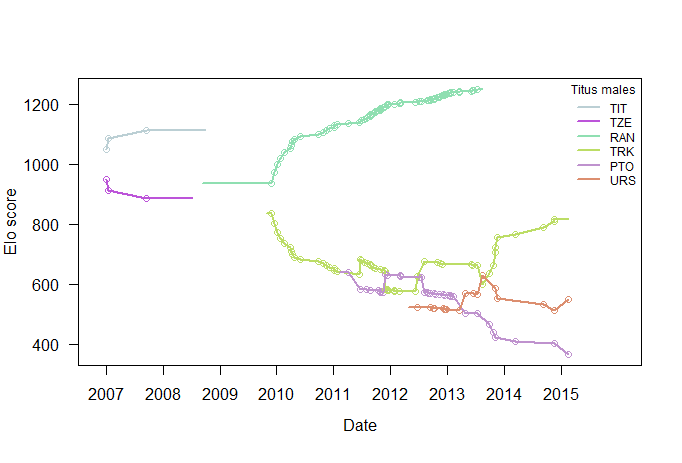

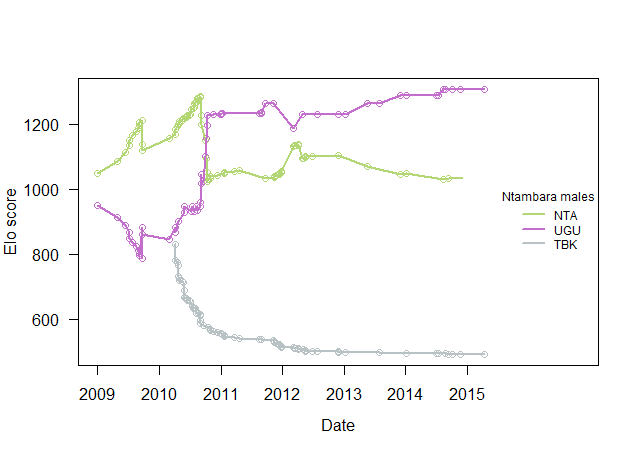

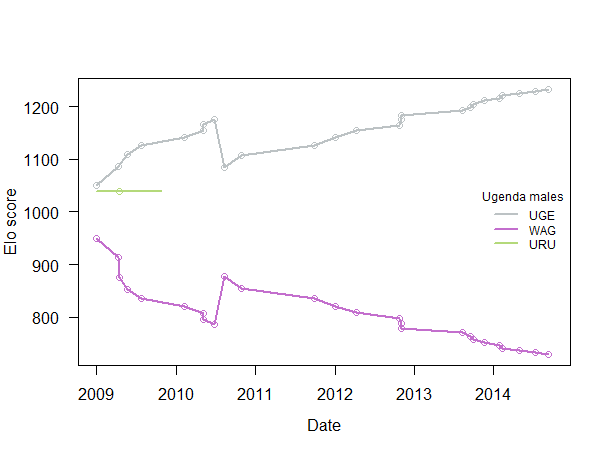

**Supp. Figure 1.** Dominance hierarchy of adult males (≤12 years) in each multi-male group with a study orphan between 2003 and 2016 established through Elo-ratings. Each colored line represents the change in Elo-rating of an adult male and each dot represents an interaction (displacement or avoidance) between two adult males. Beetsme group split into Kuryama and Titus group, and Shinda group split into Ntambara and Ugenda group, explaining the few interaction points during a period of frequent fission-fusion events in Beetsme and Shinda group before the final group split.

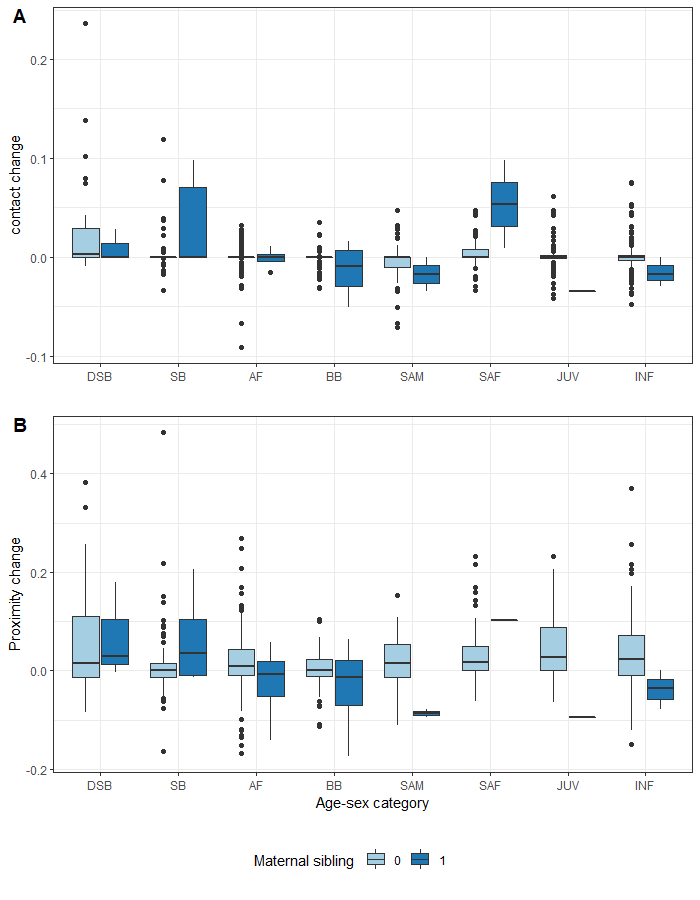

**Supp. Figure 2.** Change in relationship strength between orphans and group members that are maternal siblings (dark blue) and those that are not maternal siblings (light blue) after maternal loss based on a) affiliative contact and b) proximity.

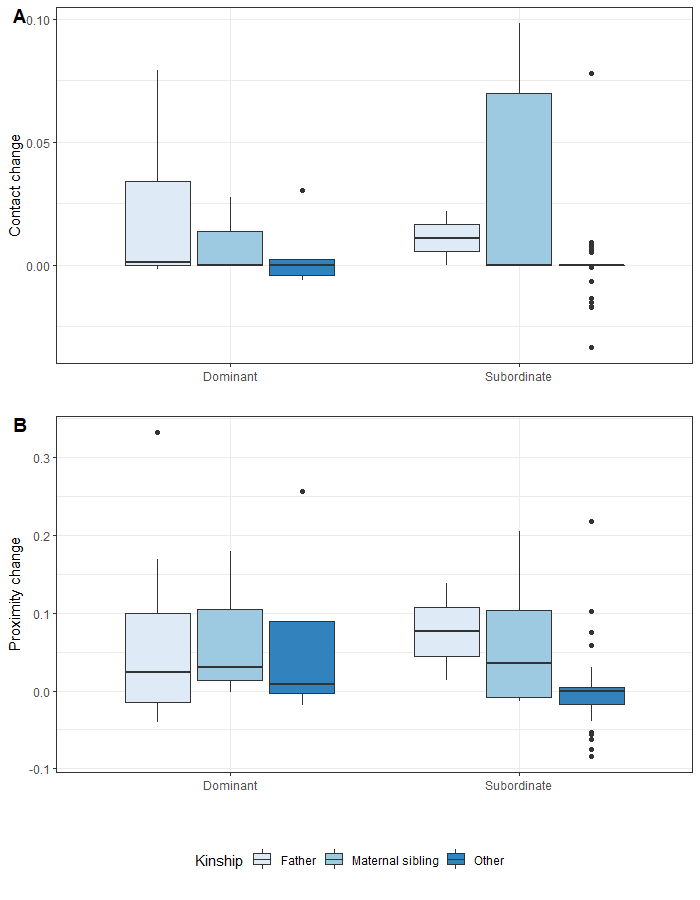

**Supp. Figure 3.** Change in relationship strength between orphans and dominant and subordinate adult male group members by kinship after maternal loss based on a) affiliative contact and b) proximity.
